# Performance of generalist hemiparasitic *Euphrasia* across a phylogenetically diverse host spectrum

**DOI:** 10.1101/2021.03.25.436816

**Authors:** Max R. Brown, Paloma G.P. Moore, Alex D. Twyford

## Abstract

- Generalist hemiparasites may attach to many different host species and experience complex parasite-host interactions. How these parasite-host interactions impact on the fitness of hemiparasitic plants remain largely unknown.
- We used experimentally tractable eyebrights *(Euphrasia,* Orobanchaceae) to understand parasite-host interactions affecting the performance of a generalist hemiparasitic plant. Common garden experiments were carried out measuring *Euphrasia* performance across 45 diverse hosts and in different parasite-host combinations.
- We showed that variation in hemiparasite performance could be attributed mainly to host species and host phylogenetic relationships (λ = 0.82; 0.17-1.00 CI). When this variation in performance is broken down temporally, annual host species cause earlier flowering, and lead to poorer performance late in the season. While *Euphrasia* species typically perform similarly on a given host species, some eyebrights show more specialised parasite-host interactions.
- Our results show that generalist hemiparasites only benefit from attaching to a limited, but phylogenetically divergent, subset of hosts. The conserved responses of divergent *Euphrasia* species suggest hemiparasite performance is affected by common host attributes. However, evidence for more complex parasite-host interactions show that a generalist hemiparasite can potentially respond to individual host selection pressures and may adapt to local host communities.

## Introduction

Parasitic plants are a diverse group of c. 4,750 species of 12 separate origins that obtain water, nutrients, and carbon from other plants using a specialised feeding organ called a haustorium (Westwood *et al.*, 2010; Nickrent, 2020). The majority of parasitic plant species are hemiparasites, which feed directly from other plants but maintain their green habit and photosynthetic competency (Twyford, 2018). These hemiparasitic plants include ecosystem engineers that reduce the growth of competitively dominant taxa in grassland communities (Pywell *et al.*, 2004), and species that threaten food security and cause billions of dollars’ worth of crop losses in agricultural systems every year (Spallek *et al.*, 2013). Generalist hemiparasitic plants may have a wide host range and attach to diverse co-occurring plant species; for example, *Rhinanthus minor* has approximately 50 host species (Gibson & Watkinson, 1989). Many aspects of the host may determine parasite performance, including nitrogen content (Korell *et al.*, 2020), carbon content (Tesitel *et al.*, 2011), secondary compounds (Adler, 2000), host condition (Houehanou *et al.*, 2011), defences (including immunity; (Cameron *et al.*, 2006; Bize *et al.*, 2008)), growth rates (Hautier *et al.*, 2010), biomass (Matthies, 2017) and genotype (Rowntree *et al.*, 2011). This complexity of host factors has impeded research into hemiparasite host range evolution, with a particular challenge being that many of these variables are confounded, and co-vary depending on the host species.

The fitness of generalist hemiparasites has traditionally been associated with host plant functional groups such as legumes, grasses, or forbs, with legumes often thought to be the best hosts (Yeo, 1964; Matthies, 1996). However, an increasing number of common garden studies have shown substantial variation in host quality within functional groups, suggesting functional group alone may not be a good predictor of host quality (Rowntree *et al.*, 2014; Matthies, 2017). Instead of functional group, many other factors, either alone, or in conjunction, could be hypothesised to explain hemiparasite performance. As some functional groups are monophyletic clades such as grasses (Poaceae), while some are paraphyletic groups such as forbs, hemiparasite performance may be better predicted by host phylogeny rather than functional group. Here, we may expect some host clades to possess attributes such as weak defences against parasites (Cameron *et al.*, 2006), or branched root architecture with many opportunities for haustorial connections (Roumet *et al.*, 2006), that confer higher parasite growth. Alternatively (or in addition), hemiparasite performance is also likely to be affected by other host attributes, for example annual or perennial life history strategies, which may have different resource accessibility (Garnier, 1992) or relative carbon and nitrogen content (Garnier & Vancaeyzeele, 1994). Finally, many theoretical models of parasitism predict that complex parasite-host interactions will arise in heterogeneous environments with variable host abundance and a mix of different host genotypes (Gandon, 2002). Such parasite-host interactions may be hypothesised to be of limited importance in facultative generalist hemiparasitic plants, where selection for host specialisation may be expected to be weak. However, growth experiments using hemiparasitic *Rhinanthus* have detected interactions between combinations of host genotype, parasite species and parasite population (Mutikainen *et al.*, 2000; Rowntree *et al.*, 2011). Such interactions are also known to be important in the obligate hemiparasitic plant *Striga*, where specific parasite-population interactions affect parasite development (Huang *et al.*, 2012). As such, parasite-host interactions may be predicted to play an important but largely overlooked role in generalist hemiparasite evolution.

Previous common garden experiments have shown substantial variation in the benefit that different hosts confer to a hemiparasite. These differences have mainly been measured as biomass or height of the hemiparasite compared to plants without a host, or between “good” and “bad” hosts (Yeo, 1964; Seel & Press, 1993; Cameron *et al.*, 2008). Few studies have tried to break down host benefits over time (Atsatt & Strong, 1970; Matthies, 1995), which may be important in natural systems with ephemeral resources and seasonal constraints, or looked at traits closely linked to fitness such as survival. Moreover, very few studies have used sufficient host replication to tease apart the general properties of host groups that influence performance. The experiments that have tested the widest range of hosts include (Matthies, 2017), who used *Melampyrum pratense* on 27 host species (Seel *et al.*, 1993) and (Rowntree *et al.*, 2014) who grew *Rhinanthus minor* on 11 host species, and (Hautier *et al.*, 2010) who used *R. alectorolophus* grown on nine host species. It is clear from these studies that as more host species are used, a wider range of hemiparasite responses, and more complex set of outcomes, will be observed. However, this variation in hemiparasite performance across many different hosts can also be leveraged to understand more general patterns, and to make direct links between how different types of host species shape the performance of hemiparasites.

Here, we use facultative generalist hemiparasitic eyebrights *(Euphrasia*, Orobanchaceae) to investigate the host attributes that determine parasite performance. This genus is an ideal model for studying hemiparasite-host interactions as they are small in size and easy to cultivate with a rapid annual lifecycle (Brown *et al.*, 2020), and species co-occur with diverse hosts in different habitats (Metherell & Rumsey, 2018). We consider multiple aspects of *Euphrasia* performance, including survival and reproduction through the year, and aim to quantify hemiparasite performance in response to many different host species. Specifically, we ask: (1) how does *Euphrasia* perform across its diverse host range and on non-hosts? (2) Do host attributes such as functional group, life history, or relatedness (phylogeny) impact on the survival and performance of hemiparasitic *Euphrasia?* (3) Do different *Euphrasia* species perform similarly with a given host species, or does reproductive success vary depending on the combination of host and parasite species (hereafter hemiparasite­ host interactions)? Our aim is to understand the potentially complex responses of a generalist hemiparasite to diverse host attributes.

## Methods

### Plant material, cultivation and trait measurements

We investigated hemiparasite-dependent host performance in two common garden experiments. Experimental 1 aimed to understand the performance of *Euphrasia* across a phylogenetic diverse spread of plant species with a range of relevant attributes such as annual and perennial life history strategies. For this experiment, we focused on a single species, *Euphrasia arctica*, due to its widespread distribution in Britain, where it mainly occupies mixed grassland habitats (Metherell & Rumsey, 2018; Becher *et al.*, 2020). We used forty-five diverse vascular plant species, including known hosts and suspected non-hosts (SI Appendix Table S1). Experiment 2 was designed to detect potential hemiparasite-host interactions using six populations from four different species of *Euphrasia* and thirteen species of hosts (SI Appendix Table S2, S3). Two diploid species (*E. anglica, E. vigursii)* and two tetraploid species (*E. micrantha, E. tetraquetra)* of *Euphrasia* were chosen to represent the diversity of the genus in Britain.

For both experiments, we used wild-collected open-pollinated seeds of *Euphrasia* (SI Appendix Table S2). Single *Euphrasia* seeds were sown in individual 9cm pots filled with Sylvamix 1 compost. Pots were placed outside at the Royal Botanical Garden Edinburgh (RBGE) in December to stratify the seeds over winter. In Experiment 1, a total of 3000 *Euphrasia* seeds were sown in winter 2016, of which 1308 germinated. In Experiment 2, a total of 2880 *Euphrasia* seeds were sown in winter 2017, of which 988 germinated. Hosts were planted in seed trays early the following spring. Following *Euphrasia* germination, plants were moved to an unheated glasshouse, and a single host introduced (Brown *et al.*, 2020). Host plants were replaced if mortality occurred within two weeks of the transplant date, and subsequently pots were randomized weekly. Plants were watered when necessary to avoid them drying out (daily in the summer), and prostrate hosts were trimmed to the edge of the pots at monthly intervals to prevent them encroaching on adjacent *Euphrasia* plants.

We measured a range of traits to understand how *Euphrasia* performance is affected by host plant species (Experiment 1) and whether specialised interactions occur between *Euphrasia* and particular host species (Experiment 2). For Experiment 1 we measured date of first flowering, and then both the number of reproductive nodes and whether an individual *Euphrasia* was alive or dead every 30 days. Survival surveys began on the 30.05.17 and ran until the 30.09.17, with these referred to as time points one (May) to five (September) herein. For Experiment 2, we measured reproductive nodes only at the end of the season. Here, reproductive nodes are the count of nodes on a *Euphrasia* plant containing either a flower or fruit, with the end of season count representing a measure of total lifetime reproductive output. In both experiments, germination date and date of host introduction were also recorded. We measured normalized transplant date, which is the time lag between germination and receiving a host, scaled to difference in first transplant date. Our analyses of hemiparasite performance were subsequently run on the following traits: number of days to flower (date of flowering – germination date), survival over time (whether an individual *Euphrasia* plant was alive at one of five time points), performance over time (number of reproductive nodes on an individual *Euphrasia* at one of five time points), and end of season performance (cumulative reproductive nodes over the lifetime of an individual *Euphrasia* plant).

### Statistical analyses

#### Hemiparasite performance across diverse host species

The statistical models for Experiment 1 were designed to assess the impact of host species and their attributes on the performance of *Euphrasia arctica.* Here, performance was measured as the number of reproductive nodes. The specific host species attributes we included were functional group of host (whether woody, a fern, forb, grass, or legume) and the life history of the host species (whether annual or perennial). We also integrated a phylogenetic tree to understand if the relatedness of putative host plants impacted the performance of *Euphrasia.* The phylogeny was based on the two gene alignment of plastid *rbcL* and *matK* from (Lim *et al.*, 2014). Six sequences from three species *(Zea mays, Hordeum vulgare* and *Lagurus ovatus)* were added from NCBI, as they were not present in the original dataset. The maximum likelihood phylogeny was generated using IQ-TREE with branch support estimated using 1000 ultrafast bootstrap replicates, and using the TESTNEWMERGE flag for model selection. A constraint tree was created using the phylomatic function in the R package brranching (Chamberlain, 2019) and used to topologically constrain the phylogeny based on the APG IV phylogeny. The tree was then made ultrametric, to scale the tree distances from root to tip, prior to model-based analyses, enabling easier calculations for the phylogenetic variance.

All subsequent analyses were conducted in R version 3.6.1 (R Core Team, 2019) with all data manipulation in base R or data.table. The three *Euphrasia* traits of interest – survival, number of days to flower, and reproductive nodes of *Euphrasia* – were modelled using a Bayesian generalized linear mixed effect model approach in the MCMCglmm package (Hadfield, 2010). This approach accommodates models with complex variance structures, and effectively handles analyses incorporating a phylogenetic tree. Four models were run with different response variables corresponding to a *Euphrasia* trait: number of days to flower, survival over time, end of season performance, and performance over time. *Euphrasia* survival was modelled using the “threshold” option in MCMCglmm, which is also known as an event history analysis model (EHA). The number of days to flower and reproductive nodes (both at the end of the season, and at each time point) were modelled using a Poisson distribution.

For all models, functional group and life history of host, as well as normalized transplant date, were added as fixed effects, whilst host species and phylogenetic effects were treated as random effects. In the EHA, time point was also added as a fixed effect to model the effect of time itself on *Euphrasia* survival. Time point five was removed from the EHA, as all but two individuals were dead at this time. We parameterized the performance over time model differently. Time point and its interaction with host life history were additional fixed effects and time points one and five were removed due to lack of reproduction. We included a random effect variance structure of an interaction of time point and host species using the us() variance function in MCMCglmm which allows covariance between host and time point:

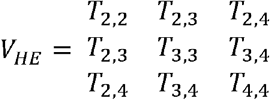

Where V_HE_ is the variance in host effect and T is the time point. The residual (V_*e*_) variance-covariance matrix allowed no covariance between time points using the MCMCglmm function idh():

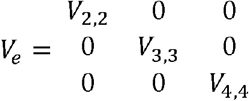

All models were run for a minimum of 130000 iterations, following a burn-in of 30000 iterations, and a thinning interval of 100. Parameter expanded priors were used to improve convergence, and effective sample sizes of focal parameters were in excess of 500 and mostly approaching 1000. Significance of categorical covariates with more than one level were determined using Wald Tests (Brown, 2019), otherwise the pMCMC value of the covariates were reported. Phylogenetic signal was calculated as the ratio of the variance of the parameter of interest to the residual variance in the model. For joint phylogenetic estimates, the posterior distributions of the phylogenetic and host species effects were summed. Significance of random effects were determined using likelihood ratio tests in the package lme4, where appropriate (Bates *et al.*, 2015). Convergence and autocorrelation of models was assessed visually by plotting the posterior distributions of the estimated parameters.

To provide a simple summary of *Euphrasia* performance comparable to the multi-host study of *Melampyrum* by Matthies (2017), we also plotted the mean performance of *E. arctica* on hosts from each functional group, including all putative hosts, and excluding likely non-hosts where *Euphrasia* produced fewer than two reproductive nodes by the end of the season.

#### Hemiparasite-host interactions

The models in Experiment 2 aimed to understand the performance of multiple *Euphrasia* species on a suite of hosts, with performance as the main response. Models were run in the R packages MCMCglmm and lme4 for significance testing of random effects. Performance was measured as the cumulative number of reproductive nodes at the end of the season, and modelled using a Poisson distribution. The fixed effects included the *Euphrasia* species, the source population (SI Appendix Table S2, and the normalized transplant date (as above). Host species and the host species interaction with *Euphrasia* species were added as single parameter random effects, as we wanted to understand the correlation in the host species effect across all *Euphrasia* species. To do this, the variances of the random effect components in our models were analysed. The correlation in host effects was calculated as:

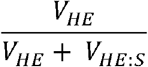

Where V_HE_ is the variance in host effects and V_HE:S_ is the variance in host species interaction with *Euphrasia* species.

All scripts for statistical analysis and figures, as well as the data used, is available at https://github.com/Euphrasiologist/euphrasiahostparasite.

## Results

### Hemiparasite performance across diverse host species

An event history analysis tracking the survival of 1308 *Euphrasia* plants through time revealed that survival was not significantly affected by host functional group (χ^2^ = 3.38, df=4, P=0.50; Fig. 1 shows legumes and grasses as examples) or host life history (χ^2^ = 0.40, df=1, P=0.53; SI Appendix Table S4). Instead, between-host effects explained 24.6% of variation in survival when accounting for phylogeny (13.4–55.4% CI, 95% Credible Intervals), with the probability of survival ranging from 0.31 when grown on heather *(Erica tetralix)* to 0.75 on cleavers *(Galium aparine).* The importance of host species was also evident from its considerable heterogeneity in effect on *Euphrasia* survival; the standard deviation of the host effects (0.57, 0.39–1.11 CI) is greater in magnitude than the fixed effects of life history (0.14, −0.25–0.61 CI) and functional group (−0.19, −1.42–0.67 CI; SI Appendix Table S4). Taken together, these results indicate host species impacts hemiparasite survival in a common garden environment, with survival being species specific rather than being influenced by host plant group (i.e. functional group, or life history).

**Figure 1.**
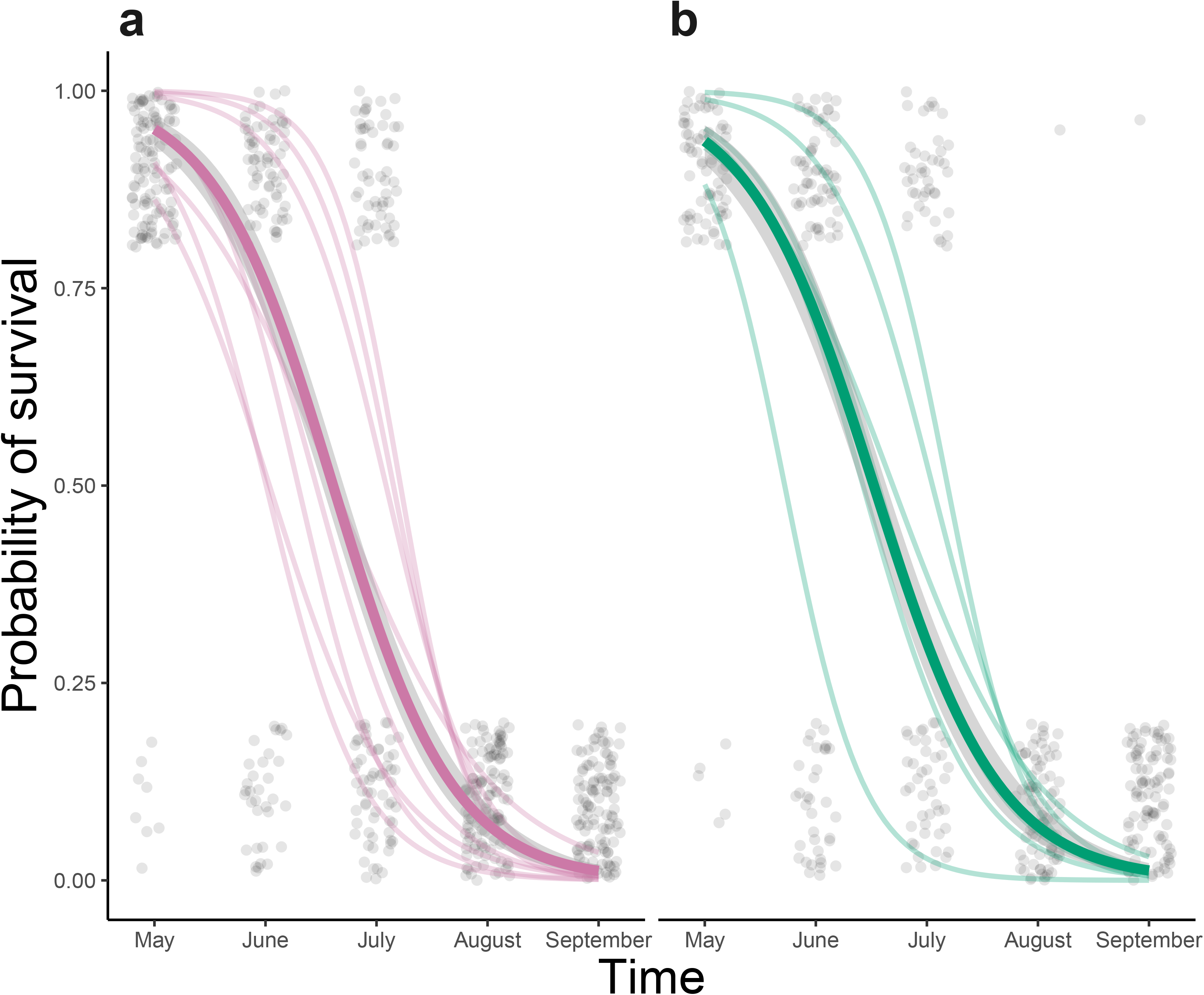
Probability of *Euphrasia arctica* surviving in a common garden experiment on 14 host species from two representative families, the Fabaceae (a) and Poaceae (b), using host species binomial regressions. Pale regressions represent individual species and bold regressions represent family level regressions. Pale grey dots are jittered raw values of an individual’s living status (binary) at each time point from earliest time point in May to the latest in August.

To understand how host species impacts on reproduction, we then tracked first flowering and reproductive success of *Euphrasia* individuals in the common garden through the growing season. The date of first flowering differed 3.5-fold across *Euphrasia* plants, with *Euphrasia* on good hosts flowering earlier (e.g. Bird’s foot trefoil, *Lotus corniculatus* = 78.0 days ± 3.5 SE, Standard Error) than those on poor hosts (e.g. maize, *Zea mays=* 129.2 days ± 5.1 SE). The difference in the number of days to flower could not be explained by host functional group (χ^2^ =2.00, df=4, P=0.73) and instead between-host effects explained 35.1% (20.0-83.5% CI) of the variation when accounting for phylogeny. Life history was marginally significant (χ^2^ =3.88, df=1, P=0.05; SI Appendix Table S5), although highly variable in its effect (77.4–101.9 days to flower CI). We found *Euphrasia* flowered earlier on annual hosts, which may be expected as annuals are a more ephemeral resource. To investigate performance over time we observed reproductive output at five time points (May­ September) throughout the season. Over this time, the effect of host functional group was non-significant (χ^2^ = 7.37, df=4, P=0.12), however host life history interacted with the September census point, with 4.7 times fewer reproductive nodes in *E. arctica* on annual hosts than perennial hosts (0.14–127 times Cl; χ^2^ = 103, df=2, P<0.001), SI Appendix Table S6). While *Euphrasia* flowered earlier on annual hosts, and therefore had the potential for a longer reproductive period, these same hosts were more likely to die earlier in the season. *Euphrasia* had consistently high reproductive success on some hosts (e.g. L *corniculatus* and *Trifolium pratense;* SI Appendix Fig. S1), however other hosts (e.g. *Cynosurus cristatus)* conferred high reproduction for *Euphrasia* earlier in the season and this then gradually declined to zero. Overall, this shows the trajectory of hemiparasite reproductive success depend on the specific host species, and their life history (SI Appendix Fig. S1).

By the end of the season, *Euphrasia* produced on average more than one reproductive node on 28 out of the 45 hosts. On average, the highest end of season performance of *Euphrasia* was observed on legumes, followed by grasses, then forbs (SI Appendix Fig. S2). However, the effects of host functional group (χ^2^ = 6.83, df=4, P=0.14, SI Appendix Table S7) and host life history (χ^2^ = 0.08, df = 1, P=0.78) were non-significant in the model based analyses. Instead, host species explained 81.8% (65.9-95.6% CI) of the variability in end of season reproductive nodes accounting for phylogeny, and phylogenetic signal was high for this trait (0.82, 0.17-1.00 CI; SI Appendix Fig S3). *Euphrasia* produced a large number of reproductive nodes only with few host species such *Lotus corniculatus* (104.5 ± 19.1 SE reproductive nodes), *Cynosurus cristatus* (53.6 ± 8.4) and the plantain *Plantago Janceolata* (35.5 ± 3.7; Fig. 2). These results highlight the importance of phylogenetic relatedness of host plant species in predicting hemiparasite performance, above host species functional group.

**Figure 2.**
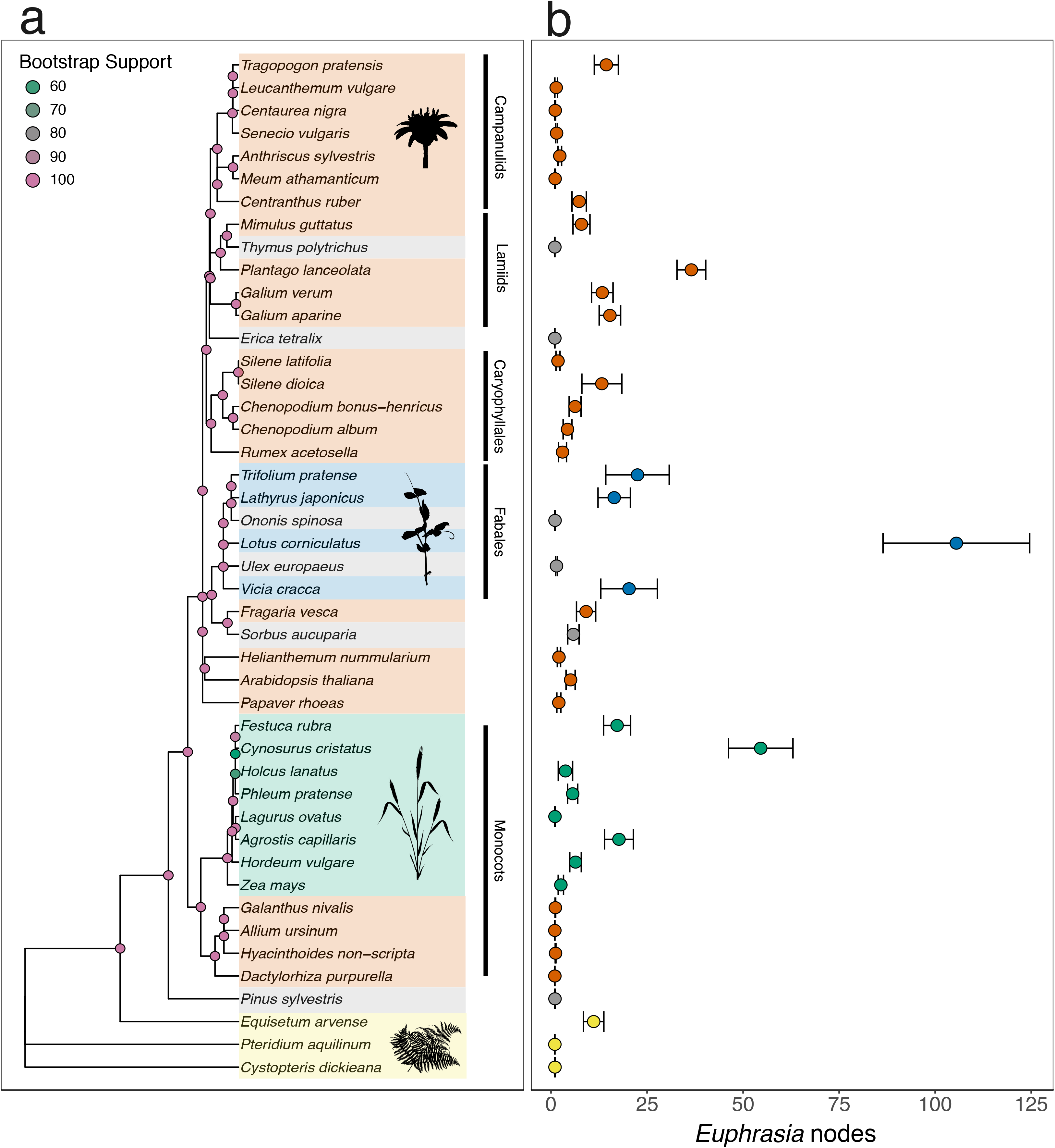
End of season performance of hemiparasitic *Euphrasia arctica* measured as cumulative reproductive nodes at the end of the season, in the context of host species and host phylogeny. (a) Maximum likelihood phylogeny of 45 hosts based on *rbcL* and *matK.* Bootstrap values are shown for each node on the phylogeny. Monocots, the two largest orders and two superorders are labelled. Host species are coloured by functional group, orange = forbs, grey = woody plants, blue = legumes, green = grasses and yellow = ferns. (b) Values are mean cumulative reproductive nodes of *Euphrasia* per species with colours corresponding to functional group of host ± one standard error. Silhouetted pictures are from phylopic.org.

### Hemiparasite-host interactions

We then tested for complex hemiparasite-host interactions, by measuring the performance of six populations from four divergent species of *Euphrasia* in a common garden using 13 hosts from different habitats (SI Appendix Tables S2, S3). A total of 635 *Euphrasia* plants survived to the end of the season. After taking into account differences between *Euphrasia* species and populations in their reproductive output (χ^2^ = 4.40, df=6, P=<0.001; SI Appendix Table S8), there was evidence for both consistent host driven differences in parasite performance, and specific hemiparasite-host interactions (Fig. 3). Host species accounted for most of the variation in reproductive nodes at the end of the season (26%; χ^2^ = 15.6, df=1, P <0.001), followed by host interacting with *Euphrasia* species (12.3%; χ^2^ = 27.1, df=1, P <0.001; SI Appendix Fig. S4). *Euphrasia* species tended to react similarly to a given host, with a 0.76 (0.37–0.93 CI) correlation in reproductive output when two hosts were picked at random (see Methods). By investigating model best linear unbiased predictors (BLUPs), we find differences in host effect are driven by L *corniculatus*, the speedwell *Veronica chamaedrys*, and sea plantain *Plantago maritima*, each of which have antagonistic interactions with different *Euphrasia* species. Moreover, two divergent species of *Euphrasia* from the same geographic location, diploid *E. vigursii* and tetraploid *E. tetraquetra*, show similar responses to the same set of hosts, with no significant interactions detected in these two species (SI Appendix Fig. SS; χ^2^ = 0.22, df=1, P=0.64). Although the dominant signal is that of conservatism of performance across *Euphrasia* species on the same host, hemiparasite-host interactions explain a significant proportion of the variation in performance.

**Figure 3.**
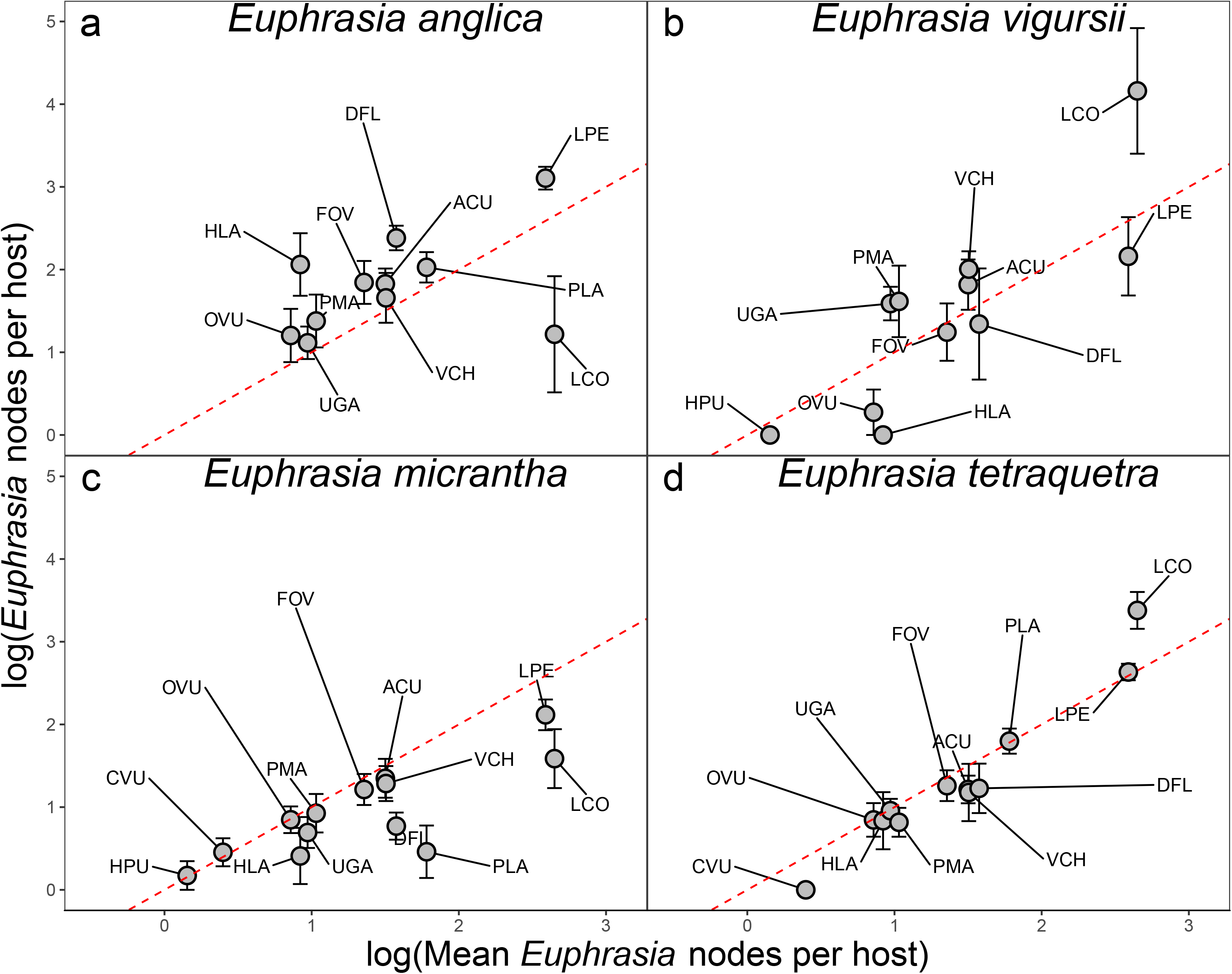
Performance of four species of *Euphrasia* on thirteen different species of host plants. Performance is measured as cumulative reproductive nodes at the end of the season. Each panel represents a unique *Euphrasia* species. The x-axis represents the number of reproductive nodes of *Euphrasia* for each host averaged across all *Euphrasia* species, while the y-axis shows reproductive nodes per *Euphrasia* species ± one standard error. Both axes are log transformed. The red dashed line graphs *y=x;* points above the line indicate elevated response to a host beyond the average, while points below the line indicate the opposite. Host species are ranked by average performance conferred to a Euphrasia species, where HPU = *Hypericum pulchrum*, CVU = *Calluna vulgaris*, HLA = *Holcus lanatus*, OVU = *Origanum vulgare*, UGA = *Ulex gallii*, PMA = *Plantago maritima*, PLA = *Plantago lanceolata*, VCH = *Veronica chamaedrys*, FOV = *Festuca ovina*, DFL= *Deschampsia flexuosa*, ACU = *Agrostis curtisii*, LPE = *Lolium perenne* and LCO = *Lotus corniculatus*.

## Discussion

We have shown that the performance of the hemiparasitic plant *Euphrasia* is determined by host attributes that impact on different aspects of survival, the initiation of reproduction, and performance through time. Our experiments used a diversity of potential host species and exposed an uneven pattern of host quality, with only a few host species providing large performance benefits. This diversity in host quality could not be directly explained by host functional group, and instead we found host quality to have strong phylogenetic signal, indicating host traits vary in a predictable way across the plant phylogeny. In addition to these observations across diverse hosts, our multi-parasite experiment uncovered evidence for both conserved and species-specific hemiparasite-host interactions. We discuss the implications of these findings in terms of the evolution of hemiparasite host range and host specialisation.

### Hemiparasite performance across a host range

We found considerable variation in host quality across forty-five putative host species, with only a subset providing substantial performance benefits to *Euphrasia.* This contrasts with the only other comparable large scale hemiparasite growth experiment to date, which found all 27 host species tested conferred some benefit to hemiparasitic *Melampyrum* (Matthies, 2017). This difference may in part be a consequence of our experiment including a larger taxonomic range spanning hosts and likely non-hosts, or may indicate that *Euphrasia* represents a more specialised hemiparasite than *Melampyrum.* Generalist parasite species are often thought to have intermediate fitness across several hosts (Leggett *et al.*, 2013), which is the case with *Melampyrum*, while *Euphrasia* performs comparatively poorly on all but a small number of genera, such as *Lotus, Cynosurus* and *Plantago. Lagurus ovatus* (grass), *Ononis spinosa* (legume), *Thymus polytrichus* (woody) and *Leucanthemum vulgore* (forb) are all putative hosts from different functional groups that conferred little to no benefit to *Euphrasia.* While legumes are on average the best host for both *Euphrasia* and *Melampyrum*, we find grasses to be next best for *Euphrasia*, while Matthies (2017) found forbs. Such comparisons between studies must be interpreted with caution due to different measure of performance, growth conditions, and hosts tested, but clearly further experimental work investigating differential host adaptation of hemiparasitic genera are warranted.

The wide variability of host quality within functional groups suggests functional group alone does not predict hemiparasite performance. This observation may be in part be due to functional group being confounded with phylogeny, with both legumes and grasses representing strongly supported clades, while forbs are paraphyletic. Our study is the first, to our knowledge, to quantify hemiparasite performance in the context of host phylogeny. The few other studies from animals and protists that have considered host phylogeny and species traits in multi-host parasite systems have also found host phylogenetic effects to be important. For example, a study of apicomplexan parasites that infect diverse bird hosts found that host phylogeny was important in explaining variation in infection status on top of environmental and host species traits (Barrow *et al.*, 2019). In *Euphrasia*, the predictive power of host relationships indicates that host traits such as defences against parasitism (Cameron *et al.*, 2006), root architecture (Roumet *et al.*, 2006), nutrient availability and the uptake of secondary compounds (Adler, 2000), and competitive ability (Keith *et al.*, 2004) are likely to vary in predictable ways across the plant phylogeny. Our experiments however, show that there are a restricted set of highly phylogenetically divergent host species which confer high benefit to *Euphrasia* (especially *L. corniculatus*, C. *cristatus* and *P. lanceolata).* Clades containing a host that confer the greatest benefits are likely to contain other species which also benefit *Euphrasia* (e.g. *Lotus, Trifolium, Lathyrus* in the legumes and *Cynosurus, Festuca, Agrostis* in the grasses). Overall, while *Euphrasia* is a true generalist able to benefit from parasitising plants throughout the vascular plant phylogeny, it only gains major benefit from attaching to a subset of taxa. *Euphrasia* species may therefore lie in a ‘grey zone’ in between generalist and specialist parasite, as has been observed in other parasitic systems (Lievens *et al.*, 2018).

### Conservation of hemiparasite-host interactions

Our finding that hosts beneficial to one *Euphrasia* species are generally beneficial across all *Euphrasia* species reveals generally conserved hemiparasite-host interactions. This is perhaps unsurprising as hemiparasites are likely to respond in a similar way to host resources, for example performing well on perennial hosts that are large, nitrogen rich and with few defences (Seel *et al.*, 1993; Cameron *et al.*, 2006; Krasnov *et al.*, 2006). While various host attributes impact hemiparasite performance, these may only be apparent when the components of plant fitness are decomposed. For example, the importance of host life history was revealed only when viewed temporally, with peak performance of *Euphrasia* on annual hosts earlier in the season. This finding highlights the ephemeral nature of annual host plants as a resource, which may be of significance in natural communities due to the restricted availability of annual hosts later in the season (Kelly *et al.*, 1988; Zopfi, 1993). Overall, the hosts that emerged as most consistently advantageous across all four *Euphrasia* species were *Lolium perenne* and *L. corniculatus*, which fulfil many of the above criteria (Beddows, 1967; Jones & Turkington, 1986). These conserved parasite responses are notable as we used highly divergent diploid and tetraploid *Euphrasia* species (~5% nucleotide divergence, corresponding to ~8 million years divergence (Wang *et al.*, 2018; Becher *et al.*, 2020)). In contrast, host conservation in many highly specialised holoparasitic taxa, like *Orobanche*, is uncommon, with host specific ecotypes found even within the same parasite species (Thorogood *et al.*, 2009).

We do however find significant hemiparasite-host interactions and species-specific responses to some hosts, suggesting weak differential host adaptation. Support for this finding can be found in the related hemiparasite *Rhinanthus*, where parasite fitness is determined by parasite genotype, host genotype and their interactions (Rowntree *et al.*, 2011)(Mutikainen *et al.*, 2000). Host species are spatially heterogenous in their distribution and vary in abundance by habitat and geographic area, creating conditions that may allow local host adaptation. The low migration rate between *Euphrasia* populations, particularly in small flowered selfing taxa (French *et al.*, 2005; Becher *et al.*, 2020), may cause differentiation and promote local adaptation. While the drivers and tempo of local host adaptation are not understood, further investigations with many hemiparasite species combined with extensive host combinations will shed light on the nature of these interactions.

## Acknowledgements

The experimental work was performed at the Royal Botanical Garden Edinburgh, which is funded from the Scottish Government’s Rural and Environment Science and Analytical Services Division (RESAS). The research was funded by a BBSRC EASTBIO DTP studentship to M.R.B. and a Natural Environment Research Council Fellowship (NE/L011336/1) and grant (NE/R010609/1) to A.D.T. M.R.B acknowledges J. Hadfield, A. Phillimore and G. Albery for modelling help.

## Author Information

### Affiliations

Institute of Evolutionary Biology, University of Edinburgh, Edinburgh, Scotland, M.R.B and A.D.T. Royal Botanical Gardens, Edinburgh A.D.T. and P.M.

### Contributions

M.R.B and A.D.T. designed the research; M.R.B and P.M. carried out the experiment and collected the data; M.R.B. analyzed the data; M.R.B and A.D.T wrote the manuscript.

### Competing interests

The authors declare no competing interests.

### Corresponding author

Correspondence to M.R.B.

## Data availability

All analyses and data are available at the online repository https://github.com/Euphrasiologist/euphrasiahostparasite

## Notes

https://github.com/Euphrasiologist/euphrasia_host_parasite

### Competing Interest Statement

The authors have declared no competing interest.

