## Supplementary Information for "Performance of generalist hemiparasitic *Euphrasia* across a phylogenetically diverse host spectrum"

Conserved and host-specific interactions in a multi-host parasite system

Max R. Brown^1,4^ (ORCID id - https://orcid.org/0000-0003-2561-516X), Paloma G.P. Moore^2^, Alex D. Twyford^1,3^ (https://orcid.org/0000-0002-8746-6617)

Corresponding authors: Max R. Brown, Wellcome Trust Genome Campus, Hinxton, Saffron Walden CB10 1RQ

Alex D. Twyford Ashworth Laboratories, Institute of Evolutionary Biology School of Biological Sciences, University of Edinburgh, Edinburgh, EH9 3JT, UK. Telephone: 0131 6508672

**This PDF file includes:**

Supplementary text

Figures S1 to S4

Tables S1 to S8

All code for analyses is available at: <https://github.com/Euphrasiologist/euphrasia_host_parasite>

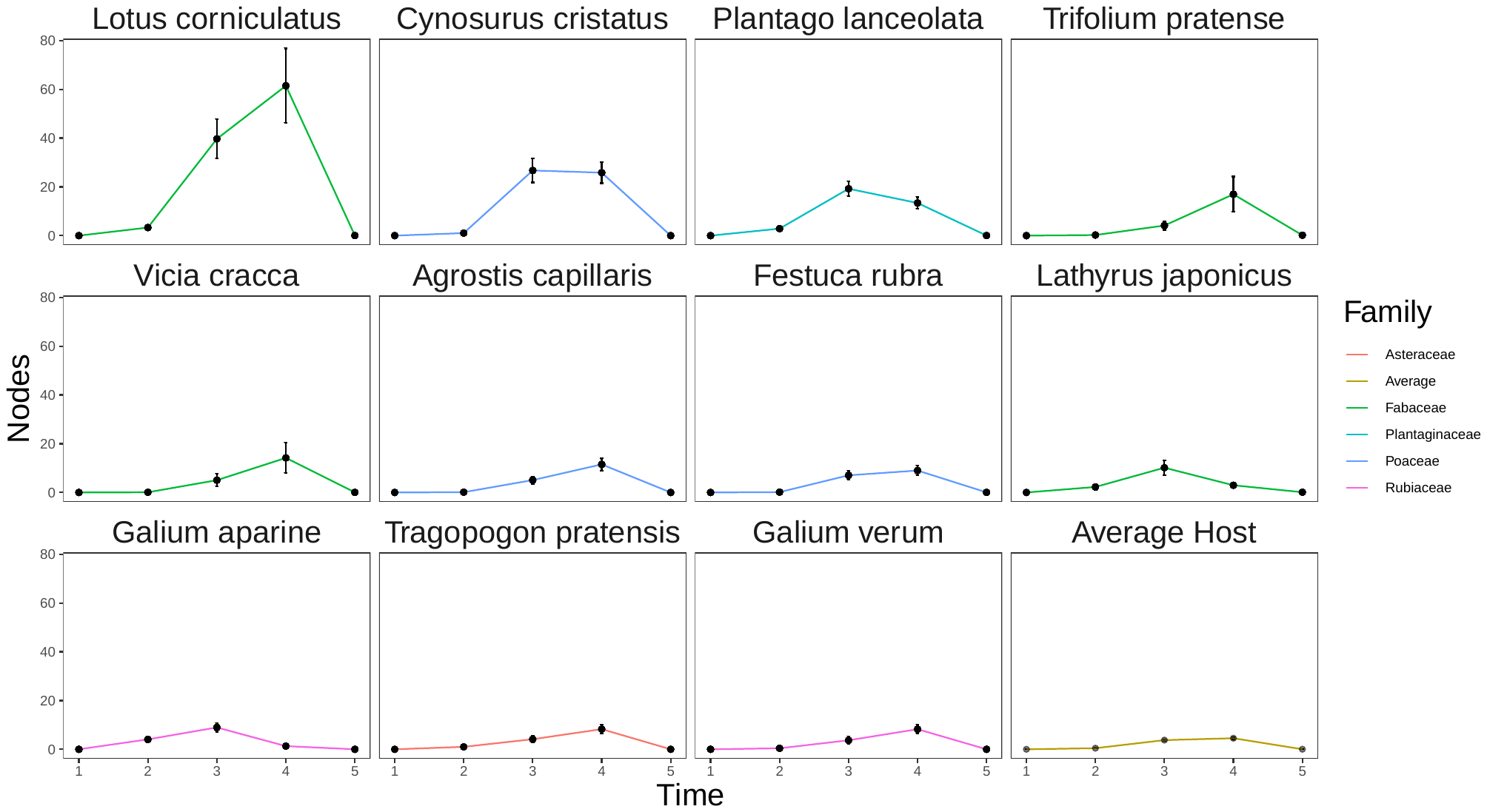

Fig. S1. *Euphrasia* reproductive output over time showing differences in reproductive trajectories, data from Experiment 1. Values represent mean reproductive nodes at a particular time point ± one standard error. Eleven species of host are shown, along with the average host where points are the mean of all hosts in the experiment.

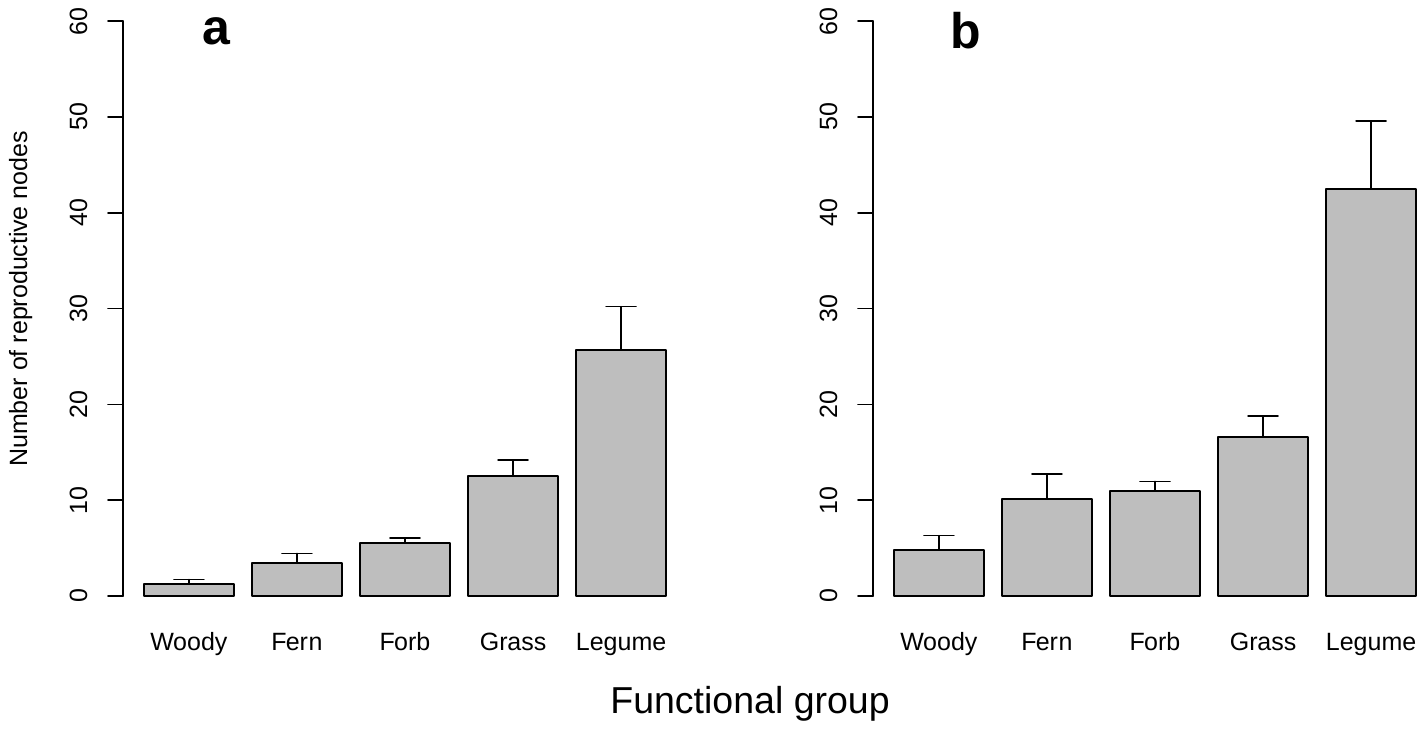

**Fig. S2.** The effect of host functional group on hemiparasitic *Euphrasia arctica* performance, measured as the mean end of season total reproductive nodes. The standard error of the mean is shown on each bar. (a) shows the performance of *E. arctica* across all host species, while (b) shows the performance of *E. arctica* on a subset of host species, excluding probable non-host species (*Allium ursinum, Anthriscus sylvestris, Centaurea nigra, Cystopteris dickieana , Dactylorhiza purpurella, Erica tetralix, Galanthus nivalis, Helianthemum nummularium, Hyacinthoides non-scripta, Lagurus ovatus, Leucanthemum vulgare, Meum athamanticum, Ononis spinosa, Papaver rhoeas, Pinus sylvestris, Pteridium aquilinum, Rumex acetosella, Senecio vulgaris, Silene latifolia, Thymus polytrichus, Ulex europaeus, Zea mays*). These host species conferred on average less than two reproductive nodes to *E. arctica* by the end of the season.

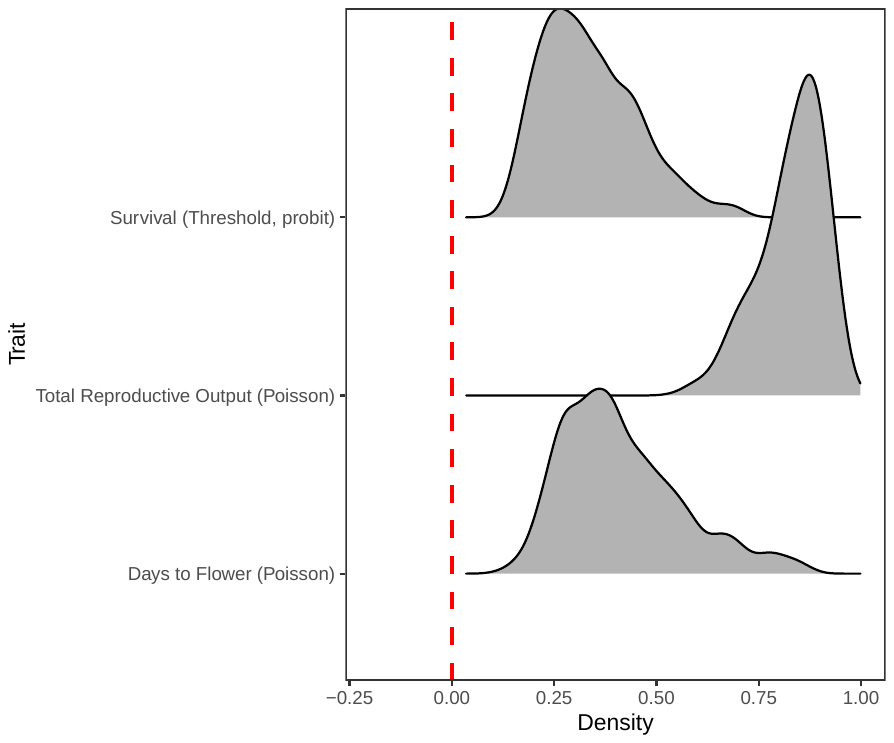

Fig. S3. Posterior distributions of the phylogenetic signal for the models from Experiment 1, where 45 different host species were grown with *Euphrasia arctica*. The distributions of phylogenetic signal are shown for three *Euphrasia* traits: survival, total reproductive output at the end of the season, and days to flower. Total reproductive output shows both the highest and least variable estimate of phylogenetic signal, however all are significant as the distributions are not overlapping zero.

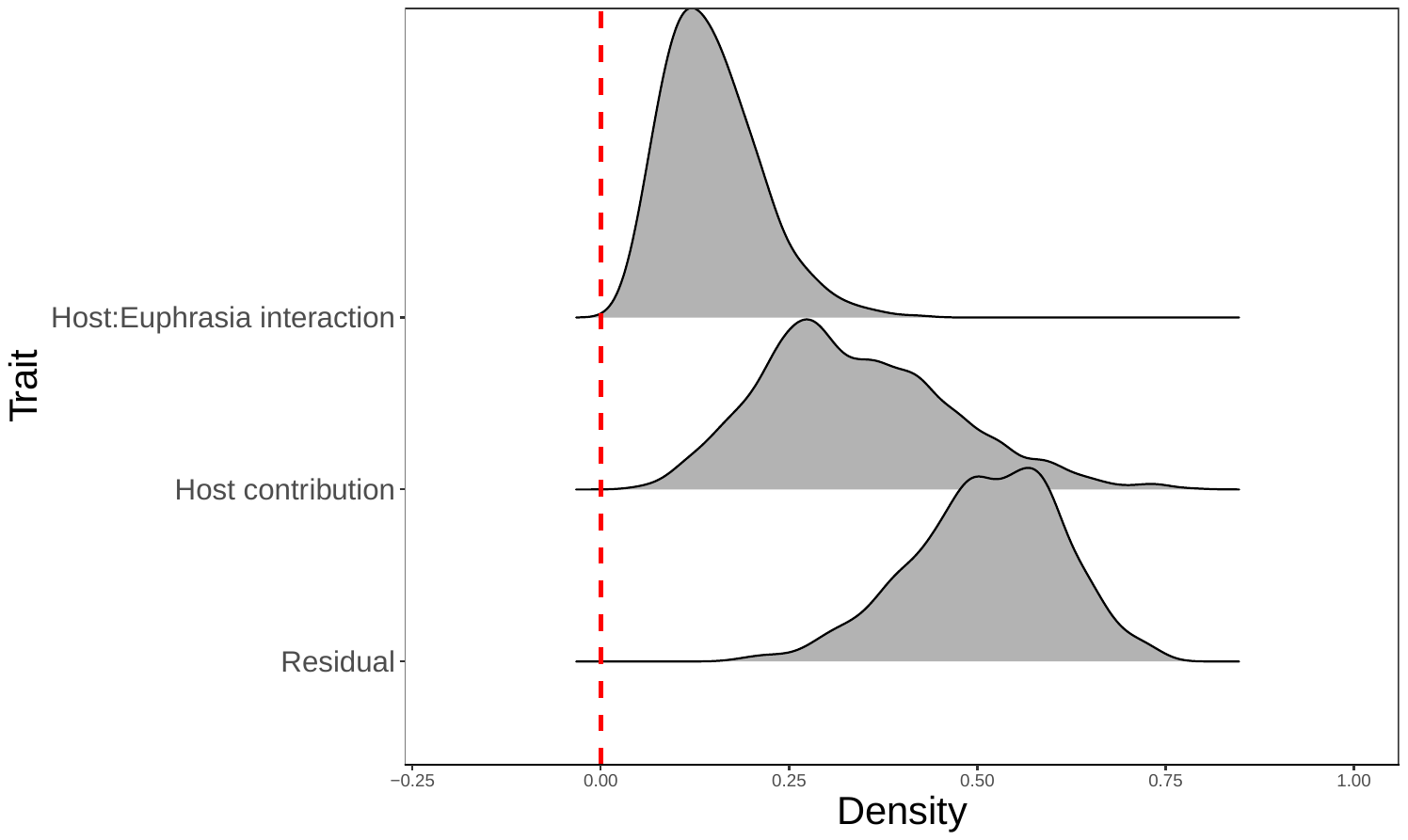

Figure S4. Posterior distribution of the variance for random effects in the model fitted for Experiment 2, where four species of *Euphrasia* were grown on thirteen different species of host. The random effects are the *Euphrasia*-host interaction, the sole effect of host species, and the residual variance. Although the residual variance is the explaining most variation, both the host-parasite interaction and hosts themselves are estimated to be significantly way from zero.

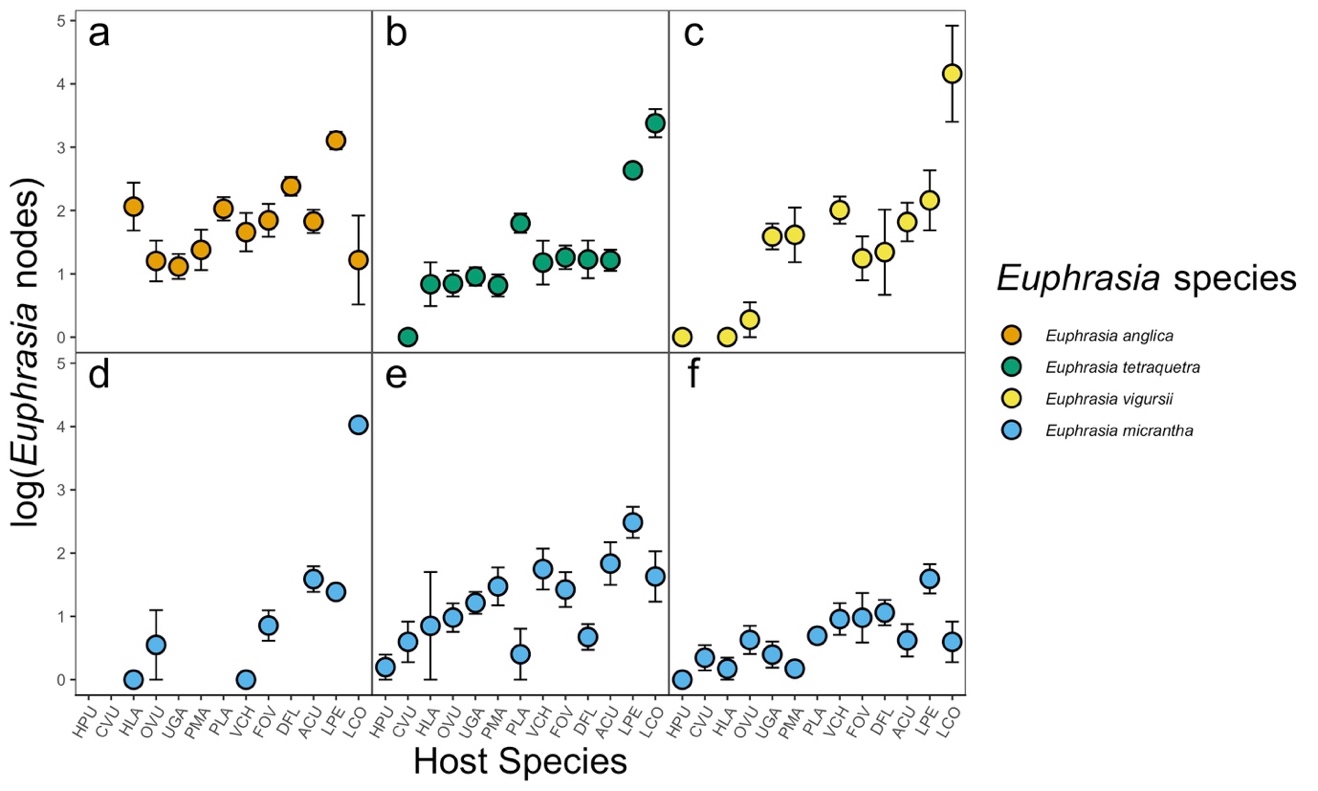

Figure S5. Performance of four species of *Euphrasia* on thirteen different species of host plants measured as cumulative reproductive nodes. Each panel represents a unique *Euphrasia* population (a = A1766, b = T1761, c = V1761, d = M1767, e = M1768, f = M1769), coloured by species. Two populations, (e) and (f) co-occur. Host species are ranked by average performance conferred to a *Euphrasia* species, where HPU = *Hypericum pulchrum*, CVU = *Calluna vulgaris*, HLA = *Holcus lanatus*, OVU = *Origanum vulgare*, UGA = *Ulex gallii,* PMA = *Plantago maritima*, PLA = *Plantago lanceolata*, VCH = *Veronica chamaedrys*, FOV = *Festuca ovina*, DFL = *Deschampsia flexuosa*, ACU = *Agrostis curtisii*, LPE = *Lolium perenne* and LCO *= Lotus corniculatus*. Y-axis values are the log of the mean cumulative reproductive nodes ± one standard error.

| **Host species** | **Authority** | **Functional group** | **Life History** | **Seed source** |
| --- | --- | --- | --- | --- |
| No host | - | - | - | - |
| *Agrostis capillaris* | L. | Grass | Perennial | Emorsgate |
| *Allium ursinum* | L. | Forb | Perennial | RBGE |
| *Anthriscus sylvestris* | [(L.) Hoffm.](http://www.theplantlist.org/tpl1.1/record/kew-2641868) | Forb | Perennial | Emorsgate |
| *Arabidopsis thaliana* | [(L.) Heynh.](http://www.theplantlist.org/tpl1.1/record/kew-2645262) | Forb | Annual | Inbred lines University of Edinburgh |
| *Centaurea nigra* | L. | Forb | Perennial | Emorsgate |
| *Centranthus ruber* | [(L.) DC.](http://www.theplantlist.org/tpl1.1/record/kew-2709046) | Forb | Perennial | Chiltern Seeds |
| *Chenopodium album* | L. | Forb | Annual | Author collections |
| *Chenopodium bonus-henricus* | L. | Forb | Perennial | Surplus seed RBGE |
| *Cynosurus cristatus* | L. | Grass | Perennial | Emorsgate |
| *Cystopteris dickeniana* | [R. Sim](http://www.theplantlist.org/tpl1.1/record/tro-26604225) | Fern | Perennial | RBGE |
| *Dactylorhiza purpurella* | [(T.Stephenson & T.A.Stephenson) Soó](http://www.theplantlist.org/tpl1.1/record/kew-55540) | Forb | Perennial | RBGE |
| *Equisetum arvense* | L. | Fern | Perennial | RBGE |
| *Erica tetralix* | L. | Woody | Perennial | RBGE |
| *Festuca rubra* | L. | Grass | Perennial | Emorsgate |
| *Fragaria vesca* | L. | Forb | Perennial | Scotia seeds |
| *Galanthus nivalis* | L. | Forb | Perennial | RBGE |
| *Galium aparine* | L. | Forb | Annual | Author collection, Upper Halliford, Surrey, Engalnd, 11/16 |
| *Galium verum* | L. | Forb | Perennial | Emorsgate |
| *Helianthemum nummularium* | [(L.) Mill.](http://www.theplantlist.org/tpl1.1/record/kew-2842625) | Forb | Perennial | Scotia seeds |
| *Holcus lanatus* | L. | Grass | Perennial | Emorsgate |
| *Hordeum vulgare* | L. | Grass | Annual | Wiggly Wigglers |
| *Hyacinthoides non-scripta* | [(L.) Chouard ex Rothm.](http://www.theplantlist.org/tpl1.1/record/kew-278557) | Forb | Perennial | RBGE |
| *Lagurus ovatus* | L. | Grass | Annual | www.wildflowershop.co.uk |
| *Lathyrus japonicus* | [Willd.](http://www.theplantlist.org/tpl1.1/record/ild-8875) | Legume | Perennial | RBGE |
| *Leucanthemum vulgare* | [(Vaill.) Lam.](http://www.theplantlist.org/tpl1.1/record/gcc-135712) | Forb | Perennial | Emorsgate |
| *Lotus corniculatus* | L. | Legume | Perennial | Emorsgate |
| *Meum athamanticum* | [Jacq.](http://www.theplantlist.org/tpl1.1/record/kew-2365193) | Forb |  | RBGE |
| *Mimulus guttatus* | [DC.](http://www.theplantlist.org/tpl1.1/record/kew-2506223) | Forb | Perennial | Author collections |
| *Ononis spinosa* | L. | Legume | Perennial | Emorsgate & Wild Flower Shop |
| *Papaver rhoeas* | L. | Forb | Annual | Emorsgate |
| *Phleum pratense* | L. | Grass | Perennial | Wild Flower Shop |
| *Pinus sylvestris* | L. | Woody | Perennial | Scotia seeds |
| *Plantago lanceolata* | L. | Forb | Perennial | Emorsgate |
| *Pteridium aquilinum* | L. (Kuhn) | Fern | Perennial | British Pteridological Society spore exchange |
| *Rumex acetosella* | L. | Forb | Perennial | Scotia seeds |
| *Senecio vulgaris* | L. | Forb | Annual | RBGE |
| *Silene dioica* | [(L.) Clairv.](http://www.theplantlist.org/tpl1.1/record/kew-2488209) | Forb | Perennial | D. Charlseworth, Univ. Edinburgh |
| *Silene latifolia* | [Poir.](http://www.theplantlist.org/tpl1.1/record/kew-2488689) | Forb | Perennial | D. Charlseworth, Univ. Edinburgh |
| *Thymus polytrichus* | [A.Kern. ex Borbás](http://www.theplantlist.org/tpl1.1/record/kew-205257) | Woody | Perennial | Emorsgate |
| *Sorbus aucuparia* | L. | Woody | Perennial | RBGE |
| *Tragopogon pratensis* | L. | Forb | Perennial | Scotia seeds |
| *Trifolium pratense* | L. | Legume | Perennial | Chiltern Seeds & Wild Flower Shop |
| *Ulex europaeus* | L. | Legume/Woody | Perennial | Tree Seed Online Ltd |
| *Vicia cracca* | L. | Legume | Perennial | Emorsgate |
| *Zea mays* | L. | Grass | Annual | Chiltern Seeds |

**Table S1:** Plant names, attributes and collection sources for host species used in Experiment 1.

| Experiment | *Euphrasia* species | Location | Grid Reference |
| --- | --- | --- | --- |
| 1 | *E.arctica* | Inverkeithing, Scotland | NT 1389 82312 |
| 2 | *E.anglica* (A1766) | Cheddar, Somerset | ST 47731 54156 |
| 2 | *E.vigursii* (V1761) | St Agnes Head, Cornwall | SW 5899 4328 |
| 2 | *E.tetraquetra* (T1761) | St Agnes Head, Cornwall | SW 5899 4328 |
| 2 | *E.micrantha* (M1767) | Borrowdale, Cumbria | NY 2468 1631 |
| 2 | *E.micrantha* (M1768) | Alness, Scotland | NH 5521 7126 |
| 2 | *E.micrantha* (M1769) | Orkney, Scotland | HY 321 055 |

**Table S2:** *Euphrasia* species collections across both experiments.

| **Host species** | **Authority** | **Source/Location** | **Plant status** |
| --- | --- | --- | --- |
| *Agrostis curtisii* | Kerguélen | Millenium Seed Bank, Kew Gardens | Seed |
| *Calluna vulgaris* | (L.) Hull | RBGE | Seed, but small plants from cuttings |
| *Deschampsia (Avenella) flexuosa* | (L.) Trin. | Chiltern Seeds | Seed |
| *Festuca ovina* | L. | Emorsgate | Seed |
| *Holcus lanatus* | L. | Emorsgate | Seed |
| *Hypericum pulchrum* | L. | Scotia Seeds | Seed |
| *Lotus corniculatus* | L. | Emorsgate | Seed |
| *Lolium perenne* | L. | Emorsgate | Seed |
| *Origanum vulgare* | L. | Emorsgate | Seed |
| *Plantago lanceolata* | L. | Emorsgate | Seed |
| *Plantago maritima* | L. | Scotia Seeds | Seed |
| *Ulex gallii* | Planch. | Millenium Seed Bank, Kew Gardens | Seed |
| *Veronica chamaedrys* | L. | Scotia Seeds | Seed |

**Table S3:** Plant names, attributes and collection sources for host species used in Experiment 2.

| **Covariates** | **Posterior mean** | **l-95% CI** | **u-95% CI** | **Effective sample size** | **pMCMC** |
| --- | --- | --- | --- | --- | --- |
| **(Intercept)** | 3.0348 | 1.8630 | 4.1519 | 1000 | **<0.001** |
| **Time** | -1.0533 | -1.1164 | -0.9912 | 1000 | **<0.001** |
| **AnnPerAnn** | 0.1390 | -0.2489 | 0.6076 | 1000 | 0.5300 |
| **Normalized transplant date** | -0.0164 | -0.0213 | -0.0117 | 1000 | **<0.001** |
| **Functional_groupFern** | -0.2583 | -1.5117 | 1.0171 | 1000 | 0.6520 |
| **Functional_groupForb** | -0.3076 | -0.9687 | 0.3844 | 1000 | 0.3700 |
| **Functional_groupLegume** | -0.0828 | -1.0457 | 0.7646 | 1000 | 0.8500 |
| **Functional_groupWoody** | -0.6675 | -1.4986 | 0.1819 | 1000 | 0.0980 |

**Table S4:** Model output from MCMCglmm for the event history analysis (survival) model in Experiment 1. The intercept represents the latent probit estimate of mean *Euphrasia* survival on a perennial grass transplanted at the earliest date, measured at the first time point. The posterior means are reported along with the lower and upper 95% credible intervals as well as the effective sample size and p-value for the effect (pMCMC).

| **Covariates** | **Posterior mean** | **l-95% CI** | **u-95% CI** | **Effective sample size** | **pMCMC** |
| --- | --- | --- | --- | --- | --- |
| **(Intercept)** | 4.6197 | 4.1765 | 5.0536 | 1000 | **<0.001** |
| **AnnPerAnn** | -0.1380 | -0.2703 | 0.0043 | 1188 | 0.0560 |
| **Functional_groupFern** | -0.1127 | -0.5410 | 0.3556 | 1000 | 0.6000 |
| **Functional_groupForb** | -0.0879 | -0.3087 | 0.1793 | 1106 | 0.3780 |
| **Functional_groupLegume** | -0.0650 | -0.3307 | 0.3032 | 860.9 | 0.6160 |
| **Functional_groupWoody** | 0.0991 | -0.2964 | 0.4466 | 1000 | 0.5520 |
| **Normalized transplant date** | 0.0034 | 0.0008 | 0.0060 | 1000 | **0.0160** |

**Table S5:** Model output from MCMCglmm for the days to flower model in Experiment 1. The intercept represents the log of the mean days to flower since germination of *Euphrasia* on a perennial grass transplanted at the earliest date. The posterior means are reported along with the lower and upper 95% credible intervals as well as the effective sample size and p-value for the effect (pMCMC).

| **Covariates** | **Posterior mean** | **l-95% CI** | **u-95% CI** | **Effective sample size** | **pMCMC** |
| --- | --- | --- | --- | --- | --- |
| **(Intercept)** | -4.1298 | -17.0773 | 5.4805 | 550 | 0.3420 |
| **Time3** | 2.3713 | 1.5862 | 3.2031 | 773.2 | **<0.001** |
| **Time4** | 3.0630 | 2.1378 | 3.9166 | 1000 | **<0.001** |
| **AnnPerAnn** | 0.7872 | -1.2385 | 2.8500 | 1000 | 0.4460 |
| **Functional_groupFern** | -4.3612 | -16.8977 | 6.6709 | 789.8 | 0.3960 |
| **Functional_groupForb** | -2.3178 | -9.4309 | 3.7584 | 793.8 | 0.4420 |
| **Functional_groupLegume** | -2.3657 | -10.7235 | 5.1473 | 756.9 | 0.5760 |
| **Functional_groupWoody** | -7.6673 | -15.5032 | -1.0839 | 549.4 | **0.0180** |
| **Normalized transplant date** | -0.0760 | -0.0919 | -0.0625 | 1000 | **<0.001** |
| **Time3:AnnPerAnn** | -0.9448 | -2.0965 | 0.1002 | 1000 | 0.0920 |
| **Time4:AnnPerAnn** | -2.3383 | -3.6057 | -0.8897 | 1000 | **0.0040** |

**Table S6:** Model output from MCMCglmm for the number of reproductive nodes over time model in Experiment 1. The intercept represents log of the mean number of reproductive nodes of *Euphrasia* on a perennial grass transplanted at the earliest date, measured at the first time point. The posterior means are reported along with the lower and upper 95% credible intervals as well as the effective sample size and p-value for the effect (pMCMC).

| **Covariates** | **Posterior mean** | **l-95% CI** | **u-95% CI** | **Effective sample size** | **pMCMC** |
| --- | --- | --- | --- | --- | --- |
| **(Intercept)** | -0.4637 | -9.8823 | 9.4058 | 1093 | 0.9240 |
| **AnnPerAnn** | -0.3610 | -2.9028 | 2.1730 | 886.5 | 0.7720 |
| **Functional_groupFern** | -3.6600 | -15.1134 | 6.8501 | 1000 | 0.4660 |
| **Functional_groupForb** | -2.9965 | -8.8016 | 2.1653 | 1097 | 0.2340 |
| **Functional_groupLegume** | -2.0488 | -9.1675 | 4.6899 | 1000 | 0.5500 |
| **Functional_groupWoody** | -7.5786 | -14.1020 | -1.0165 | 633.3 | **0.0100** |
| **Normalized transplant date** | -0.0762 | -0.0945 | -0.0570 | 1000 | **<0.001** |

**Table S7:** Model output from MCMCglmm for the cumulative reproductive nodes at the end of the season model in Experiment 1. The intercept represents the log of the mean cumulative reproductive nodes at the end of the season of Euphrasia on a perennial grass transplanted at the earliest date. The posterior means are reported along with the lower and upper 95% credible intervals as well as the effective sample size and p-value for the effect (pMCMC).

|  | **Posterior mean** | **l-95% CI** | **u-95% CI** | **Effective sample size** | **pMCMC** |
| --- | --- | --- | --- | --- | --- |
| **(Intercept)** | 1.7842 | 1.2210 | 2.2714 | 787.7 | **0.0010** |
| ***Euphrasia micrantha*** | -1.2795 | -1.7479 | -0.8284 | 1000 | **0.0010** |
| ***Euphrasia tetraquetra*** | -0.3702 | -0.8160 | -0.0076 | 873.2 | 0.0620 |
| ***Euphrasia vigursii*** | -0.2457 | -0.7758 | 0.2138 | 1000 | 0.3340 |
| **Population: M1767** | 0.3269 | -0.2098 | 0.9299 | 846.7 | 0.2760 |
| **Population: M1768** | 0.7931 | 0.4788 | 1.0699 | 1000 | **0.0010** |
| **Normalized transplant date** | 0.0059 | -0.0084 | 0.0237 | 1208 | 0.4820 |

**Table S8:** Model output from MCMCglmm for the number of cumulative reproductive nodes of Euphrasia individuals at the end of the season from Experiment 2. The intercept represents log of the mean cumulative number of reproductive nodes of *Euphrasia anglica*, population A1766, on a host that was transplanted at the earliest date. The posterior means are reported along with the lower and upper 95% credible intervals as well as the effective sample size and p-value for the effect (pMCMC).
